# DENTIST – using long reads for closing assembly gaps at high accuracy

**DOI:** 10.1101/2021.02.26.432990

**Authors:** Arne Ludwig, Martin Pippel, Gene Myers, Michael Hiller

## Abstract

**Background:** Long sequencing reads allow increasing contiguity and completeness of fragmented, short-read based genome assemblies by closing assembly gaps, ideally at high accuracy. While several gap closing methods have been developed, these methods often close an assembly gap with sequence that does not accurately represent the true sequence.

**Findings:** Here, we present DENTIST, a sensitive, highly accurate and automated pipeline method to close gaps in short-read assemblies with long error-prone reads. DENTIST comprehensively determines repetitive assembly regions to identify reliable and unambiguous alignments of long reads to the correct loci, integrates a consensus sequence computation step to obtain a high base accuracy for the inserted sequence, and validates the accuracy of closed gaps. Unlike previous benchmarks, we generated test assemblies that have gaps at the exact positions where real short-read assemblies have gaps. Generating such realistic benchmarks for *Drosophila* (134 Mb genome), *Arabidopsis* (119 Mb), hummingbird (1 Gb) and human (3 Gb) and using simulated or real PacBio continuous long reads, we show that DENTIST consistently achieves a substantially higher accuracy compared to previous methods, while having a similar sensitivity.

**Conclusion:** DENTIST provides an accurate approach to improve the contiguity and completeness of fragmented assemblies with long reads. DENTIST’s source code including a Snakemake workflow, conda package and Docker container is available at https://github.com/a-ludi/dentist. All test assemblies as a resource for future benchmarking are at https://bds.mpi-cbg.de/hillerlab/DENTIST/.

## Introduction

The quality of a genome assembly has an important impact on the quality of any downstream genomic analysis. High contiguity, completeness and accuracy of genome assemblies are fundamental to (i) comprehensively annotating genes, their promoters and other functional genomic elements, (ii) understanding genomic repeat content and organization, (iii) performing phylogenomic analysis, (iv) linking phenotypic differences to genomic differences, (v) mapping transcriptomic or epigenetic read data to an assembly, and ultimately using genomes for evolutionary and biomedical studies [1-3]. However, assembly of complex genomes is a challenging task, because sequencing reads are much shorter than chromosomes and large parts of eukaryotic genomes consist of repetitive regions [4]. Facilitated by advances in short-read sequencing that drastically increased throughput and sharply decreased costs, genomes of numerous species have been sequenced with short-read technologies such as Illumina instruments, illustrated by the Zoonomia and B10K project that generated new assemblies for 131 mammals and 267 birds, respectively [5, 6]. However, such reads are too short to span many repetitive regions, resulting in rather fragmented assemblies with short contigs (contiguous sequences) despite high read coverage. Scaffolding methods such as mate pair sequencing, chromosome conformation capture (Hi-C) read pairs or optical maps can order and orient contigs into long, sometimes chromosome-scale scaffolds. Nevertheless, due to a large number of assembly gaps, i.e. unknown genomic sequence between neighboring contigs, such assemblies remain inherently incomplete.

Pacific Biosciences (PacBio) or Oxford Nanopore Technologies instruments enable long sequencing reads, which can cover many kilobases and sometimes exceed one megabase in length. These reads are often longer than genomic repeats, which enables the assembly of highly-complete and highly-contiguous genomes, and are therefore increasingly used for *de novo* assembly [1-3, 7]. However, *de novo* long-read based assembly requires a high coverage. Approximately 60X is typically recommended for PacBio error-prone CLR (continuous long read) reads and ∼30-40X is typically recommended for the more expensive PacBio HiFi (high-fidelity) reads, which can be a significant cost factor. In addition to *de novo* assembly, a lower coverage of long reads can provide a more cost-efficient means to improve the contiguity (contig lengths) of existing short-read assemblies by bridging between neighboring contigs and thus closing assembly gaps. Furthermore, gap filling is also one of the final steps in state-of-the-art pipelines to generate high-quality long-read assemblies [1]. To close gaps in existing fragmented assemblies with more limited long-read coverage, several methods have been developed in the past, exemplified by PBJelly [8], FinisherSC [9], PacBio GenomicConsensus (PacBio GC, https://github.com/PacificBiosciences/GenomicConsensus/) comprising the Arrow and Quiver algorithm implemented in the variantCaller tool, LR_gapcloser [10], and TGS-Gapcloser [11]. These methods align a set of given long reads to an input short-read assembly, determine which reads span assembly gaps and close these gaps with the new sequence. However, as we demonstrate below, even for shorter and less repeat-rich genomes, these tools lack high accuracy when closing assembly gaps. Thus, while applying these tools improves assembly contiguity, they compromise the quality of the resulting assembly, which can impair downstream analyses.

Here, we developed DENTIST, a new, reliable and sensitive gap closing method that unlike previous methods achieves a high level of accuracy. Using various simulated and real data sets for genomes ranging from small (*Drosophila*) to large and complex (human), and using realistic assembly gap loci, we demonstrate that DENTIST consistently achieves the highest accuracy compared to other state-of-the-art approaches, while having a similar or better runtime and memory consumption.

## Results

### Overview of DENTIST

DENTIST implements a full gap closing pipeline (Figure 1) and was developed with the main consideration of closing assembly gaps at a very high accuracy. Conceptually, given an assembly and a set of long reads, gap closing starts by aligning the reads to the assembly to identify those that reach into or span assembly gaps. The reads are then used to infer the DNA sequence that should be used to fill these gaps.

**Figure 1:**
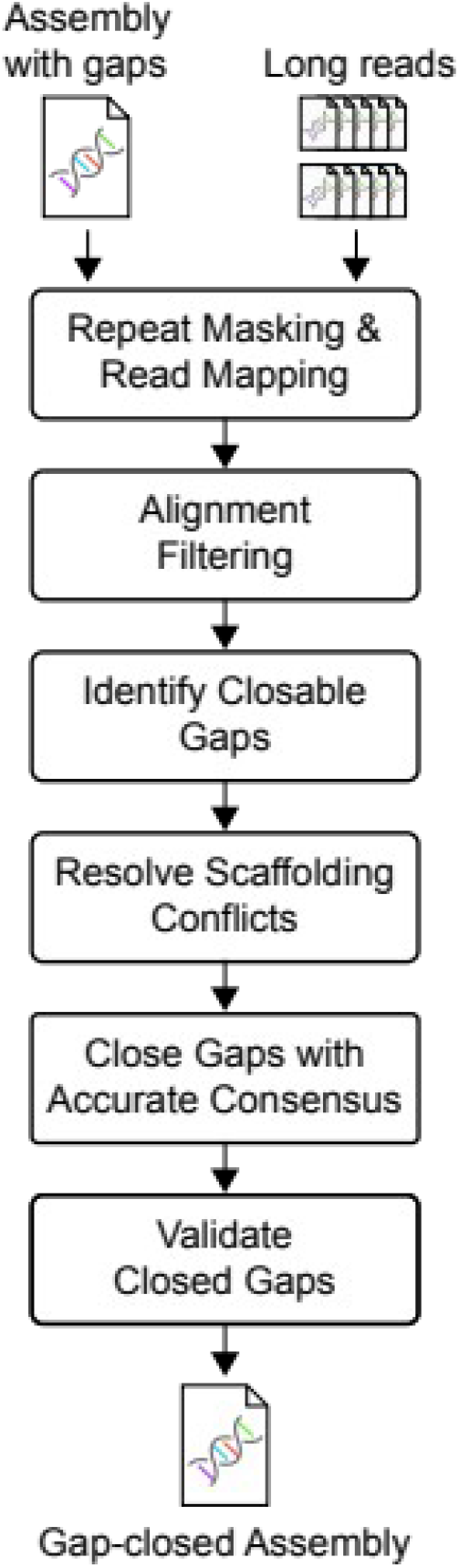
Overview of the assembly gap closing pipeline implemented in DENTIST.

DENTIST differs from previous approaches in the following key aspects. First, a key step in accurately closing gaps is to align the long reads to the correct loci in the input assembly. This is not a trivial task, since genomic repeats can lead to ambiguous read alignments and potentially assigning reads to the wrong genomic locus. Therefore, DENTIST integrates four approaches to identify repetitive regions in both the input assembly and the input long reads, and explicitly uses this repeat mask to identify reliable read alignments. Many of the remaining alignment ambiguities and conflicts are resolved using a scaffolding graph. Second, as long reads generally have a high base error rate, it is important to determine an accurate consensus sequence from the long reads that span or overlap an assembly gap before closing the gap. DENTIST employs a state-of-the-art reference-based consensus caller to generate high-quality consensus sequence for each closed assembly gap, maintaining a high base accuracy in the final assembly. Third, aiming at a high accuracy, DENTIST validates both the consensus sequence as well as the closed gap by aligning the input reads again to the gap-closed assembly, and does not perform questionable gap closures in favor of a correct result.

### Test procedure

To assess the performance of DENTIST and compare it to existing methods, we followed previous strategies to generate a “ground truth scenario” where assembly gaps were inserted into a high-quality assembly and long reads were used to close the created assembly gaps. This general strategy allows one to compare sensitivity (number of closed gaps, increase in assembly contiguity) and, since the real gap sequence is known, the accuracy of the inserted sequence.

Unlike previous comparisons, we devised a ground truth scenario, where the assembly gap sizes and loci are as realistic as possible, which is important to assess real-life performance. Previous comparisons introduced assembly gaps at random locations [8] or replaced repeats by assembly gaps of the same length [10]. By excising repeats, the latter approach results in gap flanks having often unique, non-repetitive sequence that make accurate read alignment easier. Consequently, both approaches do not reflect the complexity of assembly gap sizes and locations in reality, and the fact that contig ends often reach into the unbridgeable repeat (as opposed to ending right before it). To overcome this, we devised a procedure that uses two real assemblies of the same species. A real high-quality assembly constitutes the “ground truth” and a real fragmented short-read assembly is used to obtain realistic assembly gap locations. Specifically, we aligned the fragmented assembly to the ground truth assembly and introduced gaps into the ground truth assembly at the exact position and size where gaps occurred in the short-read assembly (Figure 2). Then we either used long reads that were sampled (simulated) from the ground truth assembly or real PacBio reads of the same species (Supplementary Table 1) to close assembly gaps and evaluate sensitivity and correctness.

**Figure 2:**
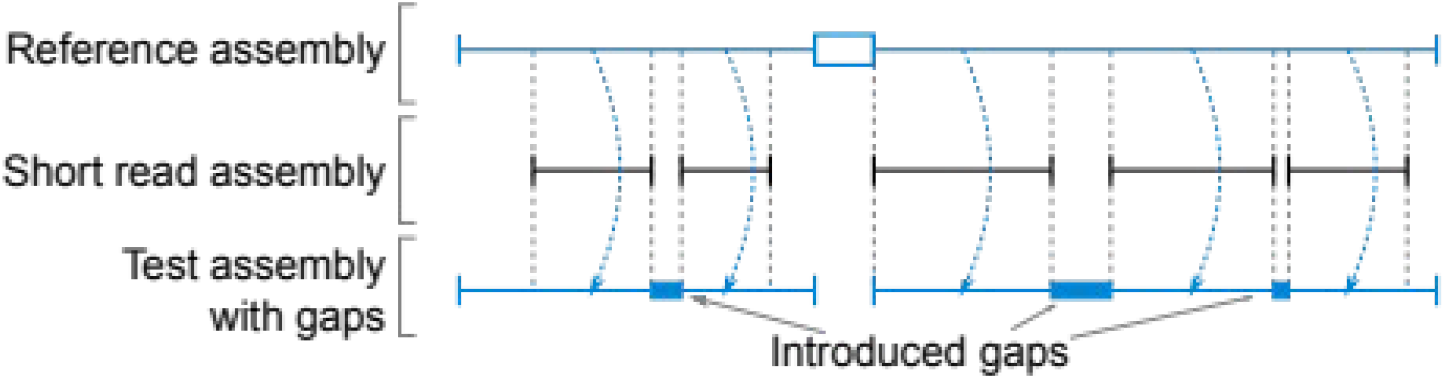
Realistic ground truth scenarios are created by “copying” gaps from a real short-read assembly to a real high-quality reference assembly. Horizontal lines represent contigs, blue boxes represent genomic regions of the reference assembly, which were replaced with assembly gaps (N’s). The white box in the reference assembly represents an assembly gap in the reference, which is not copied to the test assembly.

In the following comparison, we applied DENTIST and five other state-of-the-art methods, PBJelly [8], FinisherSC [9], PacBio GenomicConsensus (https://github.com/PacificBiosciences/GenomicConsensus), LR_gapcloser [10], and TGS-Gapcloser [11] to four species with various genome sizes and repeat content, representing the insect, plant and vertebrate lineage: the fruit fly *Drosophila melanogaster* with a 134 Mb assembly where ∼10% of the ground truth assembly was covered by DENTIST’s repeat masks and 1,340 gaps were introduced, the thale cress *Arabidopsis thaliana* (119 Mb, ∼11% repeats, 219 gaps), the hummingbird *Calypte anna* (1 GB, ∼2% repeats, 37,501 gaps), and human (3 Gb, ∼20% repeats, 5,846 gaps). Sources and details of all assemblies and reads as well as statistics of introduced gaps are listed in Supplementary Table 1. We were only able to test FinisherSC on the smallest genome (*Drosophila*), as this method failed on *Arabidopsis* with simulated reads, did not finish within two days on *Arabidopsis* with real PacBio reads and required more than 1 TB of RAM for larger genomes. Since gap and assembly properties differ between the four species, we note that the following comparisons of different methods only provide a relative performance assessment on same input dataset.

### Simulated Read Data

We first simulated PacBio reads from each ground truth assembly with a typical PacBio error profile and an average raw read length of 26.7 kb (Supplementary Table 1). For *Drosophila, Arabidopsis* and hummingbird, the gap sequence inserted by DENTIST is on average ≥99.92% identical to ground truth sequence (Table 1, Figure 3A). In comparison, the sequences inserted by LR_gapcloser, PBJelly and TGS-Gapcloser had a lower average identity to the ground truth sequence (∼87.7% for LR_gapcloser, 88.61–95.17% for PBJelly, 92.81-97.26% for TGS-Gapcloser). FinisherSC, which we could only run on the *Drosophila* dataset, achieved an identity of only 81.53% (Table 1). For the 3 Gb human genome, the average sequence identity for DENTIST was 98.70%, whereas LR_gapcloser (88.01%), PBJelly (89.25%) and TGS-Gapcloser (93.63%) achieved lower average identity values. In addition to a high average identity, DENTIST closed 46.1% (human) to 91.5% (*Drosophila*) of the gaps with a 100% accurate sequence, while other methods at best closed 17.6% (TGS-Gapcloser, 236 of 1,340 gaps for *Drosophila*) at 100% accuracy.

**Table 1:**
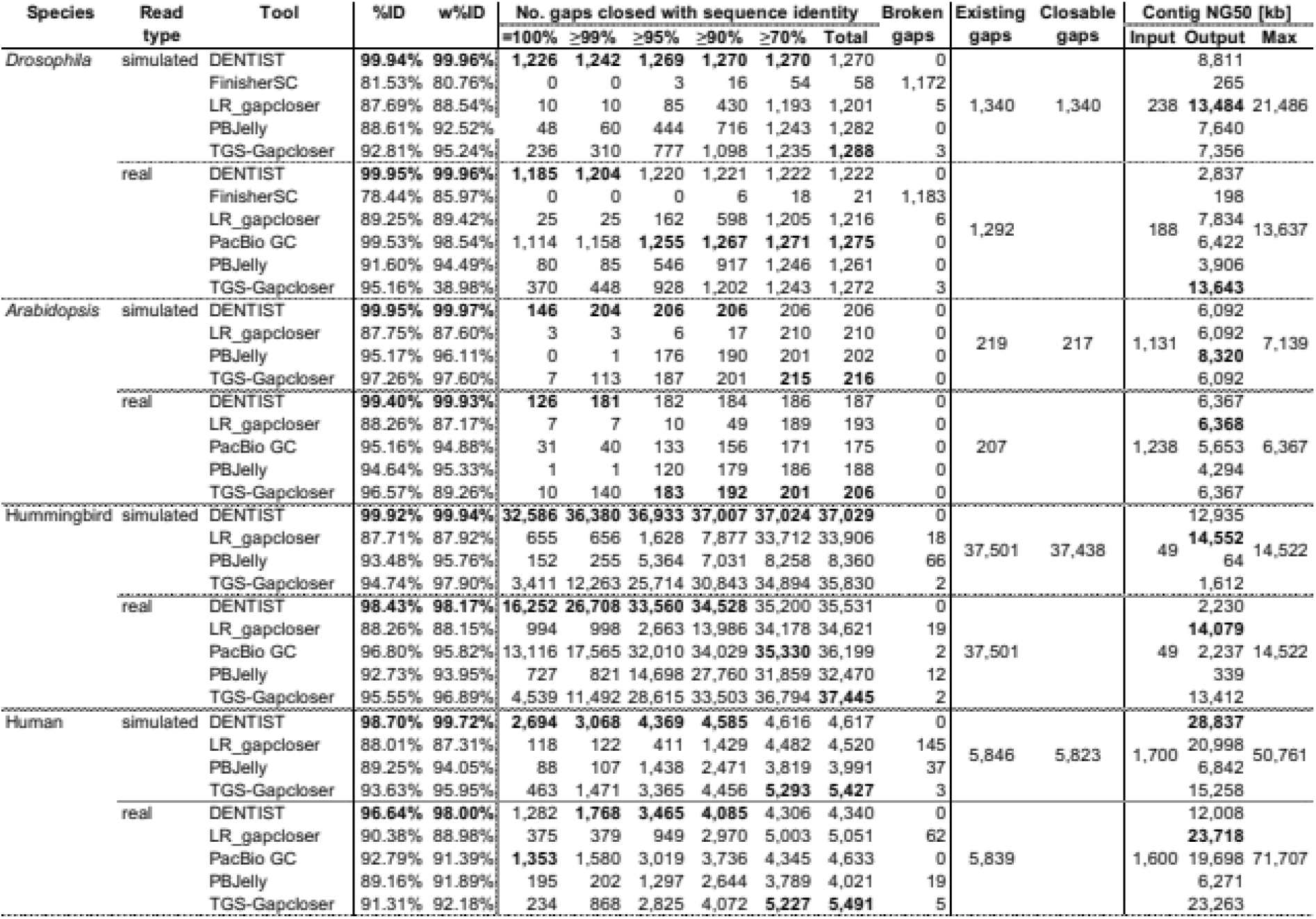
Comprehensive benchmarking of gap closing methods. We define %ID as the average across all sequence identities of closed gaps and w%ID as the average weighted by the true gap length. Gaps are broken if the flanking contigs are not adjacent anymore after gap closing, indicating a mis-assembly. For simulated reads, we label a gap as closable if there are at least three simulated reads that span the gap and at least 500 bp of its flanks. The contig NG50 value of the test (input) and output assembly is given on the right; maximum contig NG50 refers the value if all introduced gaps would be closed. Best values in each category are in bold.

**Figure 3:**
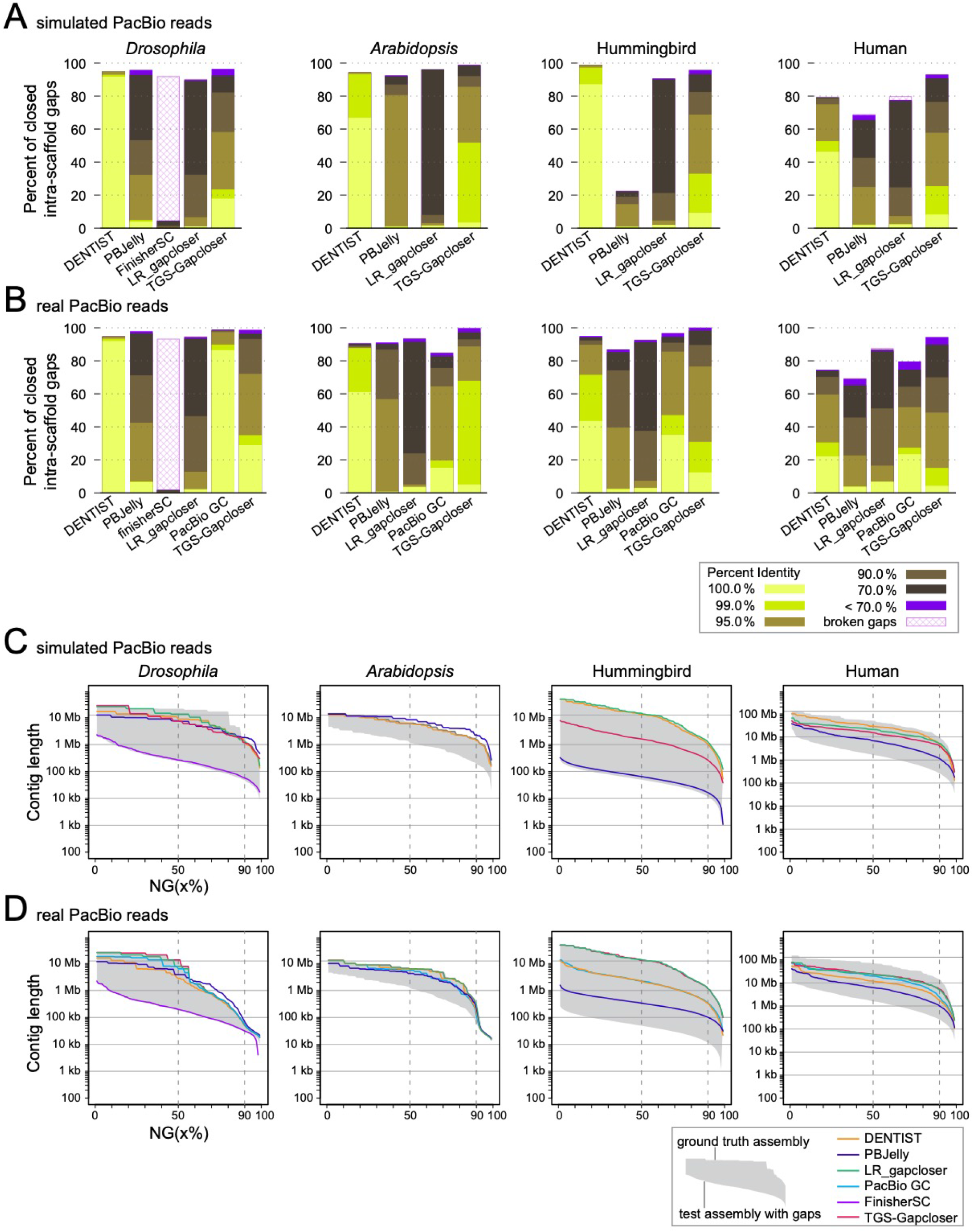
Comparison of accuracy and sensitivity of gap closing methods. (A,B) Bar charts show the percent of gaps that are closed by different methods using simulated (A) or real (B) PacBio reads and a breakdown of how identical the inserted sequence is to the ground truth sequence. (C, D) NG(x) plots show which percent of the assembly consists of contigs of a certain minimum size (Y-axis). The contig NG50 and NG90 values are indicated by vertical dashed lines. The grey area represents the NG(x) of the test assembly (contains introduced assembly gaps, lower bound) and the ground truth assembly (representing the maximally-possible contiguity if all introduced gaps are closed, upper bound). For *Drosophila* (simulated and real reads) and *Arabidopsis* (simulated reads), PBJelly shows NG(x) values higher than the values of the ground truth assembly, indicating that some gaps are “overfilled”. Due to excessive memory consumption, we could run FinisherSC only on the *Drosophila* genome and this method produced many broken gaps (likely representing mis-assemblies, as the input contigs are not adjacent anymore in the output assembly). PacBio GC requires real read data as input and is therefore only shown in panel D. For *Arabidopsis* and real PacBio reads (panel D, second column), the TGS-Gapcloser line overlaps the LR_gapcloser line and follows the upper bound.

To account for the different sizes of assembly gaps, we also calculated the average identity weighted by the size of the gap. This metric showed highly similar values (Table 1), with the exception of PBJelly and TGS-Gapcloser where the weighted average identity increased by 0.94–4.8% and 0.34–3.16%, respectively, indicating that larger gaps tended to be closed with a higher accuracy (discussed below). Nevertheless, the maximum weighted average of PBJelly (96.11% for *Arabidopsis*) and TGS-Gapcloser (97.9% for hummingbird) is lower than the accuracy of DENTIST (always ≥99.72%). In summary, consistently for all assemblies, DENTIST achieves the highest accuracy.

Comparing sensitivity, we found that all methods closed a similar percentage of gaps for *Drosophila* (89.6-96.1%, except FinisherSC with 4.3%), *Arabidopsis* (92.2-98.6%) and hummingbird (90.4-98.7%, except PBJelly with 22.3%) (Table 1, Figure 3A). TGS-Gapcloser closed the most gaps for *Drosophila* (96.1%), and *Arabidopsis* (98.6%), while DENTIST closed the most gaps for hummingbird (98.7%). For the larger human assembly, TGS-Gapcloser closed the most gaps (92.8%) followed by DENTIST (79%). To measure the increase in assembly contiguity, we computed the contig NG50 value, using the number of real bases (A,C,G,T) in the ground truth assembly as a fixed reference. This allows us also to compute the maximally possible contig NG50 that can be achieved by closing all introduced gaps in the ground truth assembly. It should be noted that contig NG50 increase is not a perfect measure of sensitivity, as the NG50 increase heavily depends on the position of closed gaps and reflects only indirectly the number of closed gaps. For example, closing a single gap between two large contigs could result in a larger NG50 increase than closing several gaps between several small contigs. Indeed, this effect is also visible in our comparisons, exemplified by the hummingbird assembly where LR_gapcloser achieved a higher contig NG50 (14.6 Mb vs. 12.9 Mb) despite DENTIST closing 3,123 more gaps (Table 1). Despite these caveats, comparing contig NG50 increase, we found LR_gapcloser achieved the highest contig NG50 for *Drosophila* and hummingbird, PBJelly achieved the highest contig NG50 for *Arabidopsis*, and DENTIST achieved the highest contig NG50 for human (Table 1). For all four species, DENTIST substantially increased the contig NG50 between 5-fold (*Arabidopsis*) and 263-fold (hummingbird). We note that PBJelly outputs an *Arabidopsis* assembly with a significantly higher contig NG50 value than what is achievable by correctly closing all introduced gaps (8.3 vs. 7.1 Mb), which indicates that some gaps may be “overfilled”. Since NG50 reduces contiguity to a single number, we also plotted which percent of the assembly is contained in contigs exceeding a certain size (Figure 3C). These NG(x) plots showed that DENTIST, PBJelly, LR_gapcloser and TGS-Gapcloser achieve similar improvements in contiguity for *Drosophila* and *Arabidopsis*, that DENTIST and LR_gapcloser achieve a higher contiguity for hummingbird, and that DENTIST achieves a higher contiguity for human. In summary, our comparisons using simulated reads show that the sensitivity of DENTIST is comparable and sometimes better than other methods and that DENTIST excels in a very high accuracy.

### Real PacBio long-read data

Since simulated reads cannot capture the full diversity of issues that are present in real PacBio reads (e.g. chimeras, low quality regions, missed adapters) and further do not encompass the whole genome but were only sampled from the ground truth contigs, we next evaluated the performance of DENTIST and other gap closing methods on the same species but using real CLR PacBio reads (Table 1). Read length, gap and ground-truth assembly statistics are listed in Supplementary Table 1. We added the PacBio GC method to the tests, which requires additional sequencing metrics (such as pulse widths and inter-pulse durations) as input and can therefore only be applied to real read data. We note that in these tests, it will be harder to achieve a very high accuracy as real reads also contain SNPs or larger haplotype variation and the ground truth assembly may contain base or assembly errors with respect to the real reads. Nevertheless, the accuracy of DENTIST is still very high with an average identity between inserted and ground truth sequence of 99.95% for *Drosophila*, 99.40% for *Arabidopsis*, 98.43% for the hummingbird and 96.64% for human (Table 1, Figure 3B). All other methods have a lower accuracy. The second best methods are PacBio GC for *Drosophila* (99.53%), hummingbird (96.80%) and human (92.79%) and TGS-Gapcloser for *Arabidopsis* (96.57%). A few examples of closed gaps are illustrated in Figure 4.

**Figure 4:**
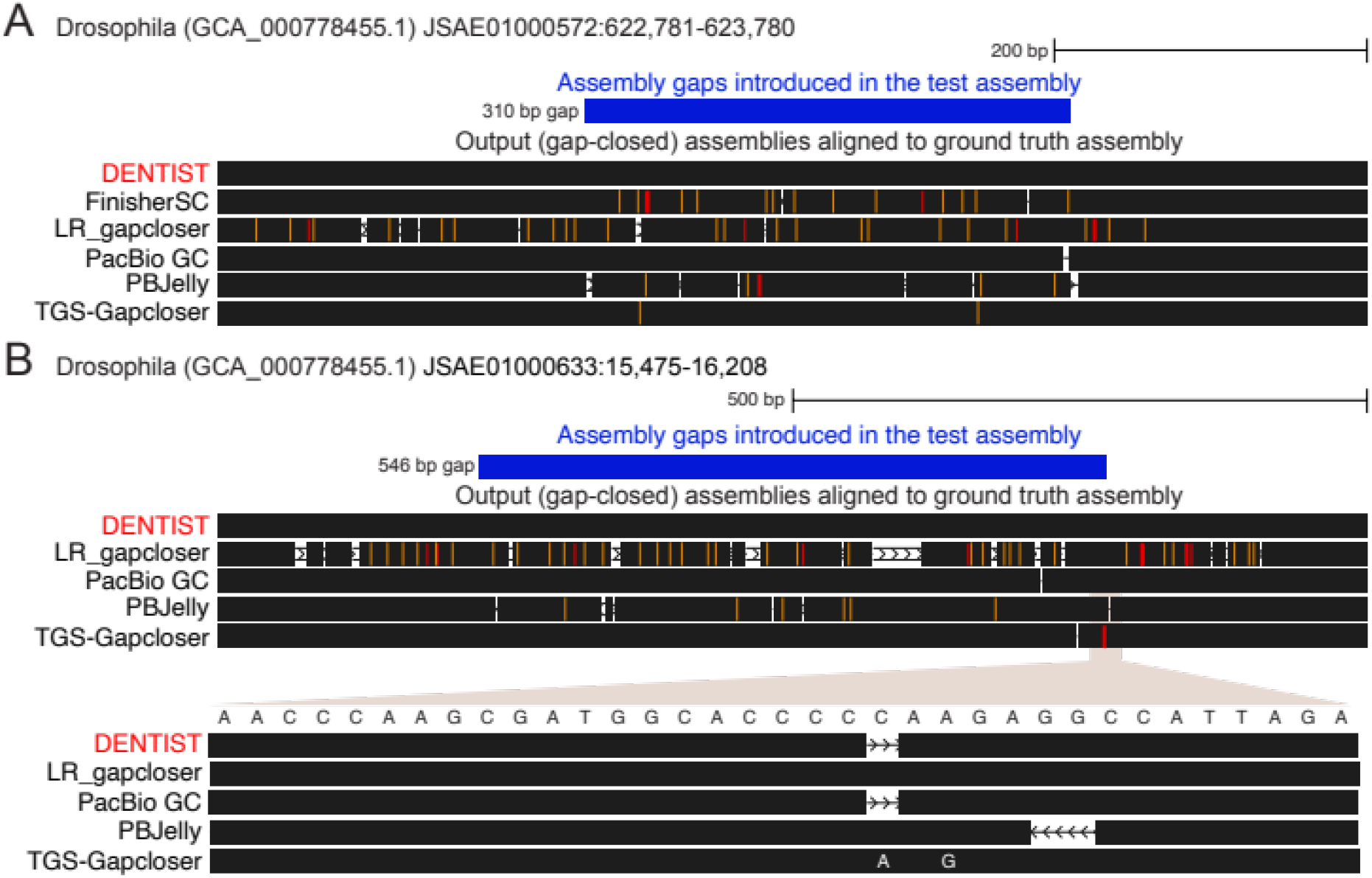
Examples of gaps in the *Drosophila* assembly that are closed with real PacBio reads. UCSC genome browser [20] visualizations show a part of the ground truth *Drosophila* assembly overlapping an assembly gap that we introduced in the test assembly (blue). The output assemblies produced by different gap closing methods are aligned to the ground truth assembly, highlighting base differences in color (deletions in white, insertions in orange, substitutions in red) and identical sequence parts in black. (A) While DENTIST closes the 310 bp gap with a sequence that is 100% identical to the ground truth, other methods introduce a few base errors (98.7% identity for TGS-Gapcloser and PacBio GC; lower for other methods). (B) DENTIST closes the 546 bp gap with a sequence that is 99.8% identical to the ground truth, while other methods have slightly lower identity values (PacBio GC 99.5%, TGS-Gapcloser 99.1%, PBJelly 96.4%, LR_gapcloser 84.2%). The inset shows that the single base error in the DENTIST output is a deletion of a ‘C’ in a short homopolymer run of C’s, and that PacBio GC makes the same mistake.

For human, DENTIST’s average identity weighted by gap size is noticeably higher than the unweighted identity (98.00% vs. 96.64%), indicating that larger gaps tend to be closed with a higher accuracy while some of the smaller gaps were closed less accurately. We confirmed this by plotting identity vs. gap size (Figure 5). Manual inspection showed that many of these less accurate cases comprise gaps <50 bp consisting of simple repeats such as homopolymer runs, for which long reads have a higher error rate [12], making it more difficult to compute an accurate consensus. Since very few incorrect bases in the consensus of a small gap have a large effect on the average identity (e.g. 19 of 20 correct bases is an identity of only 95%), the weighted average identity provides a better measure of overall sequence accuracy. Furthermore, few base errors in a small gap may not be a serious problem, since small gaps can easily be “polished” with Illumina reads in a downstream step (discussed below). A consistently higher weighted average identity was also observed for PBJelly (Table 1, Figure 5B). For TGS-Gapcloser, the weighted identity is significantly reduced for *Drosophila* (95.16% vs. 38.98%) and *Arabidopsis* (96.57% vs. 89.26%), suggesting that some large gaps were filled with a low accuracy. In summary, considering both the weighted and unweighted average identity of the inserted sequence, DENTIST achieves the highest accuracy consistently for all four assemblies also for real PacBio CLR reads.

**Figure 5:**
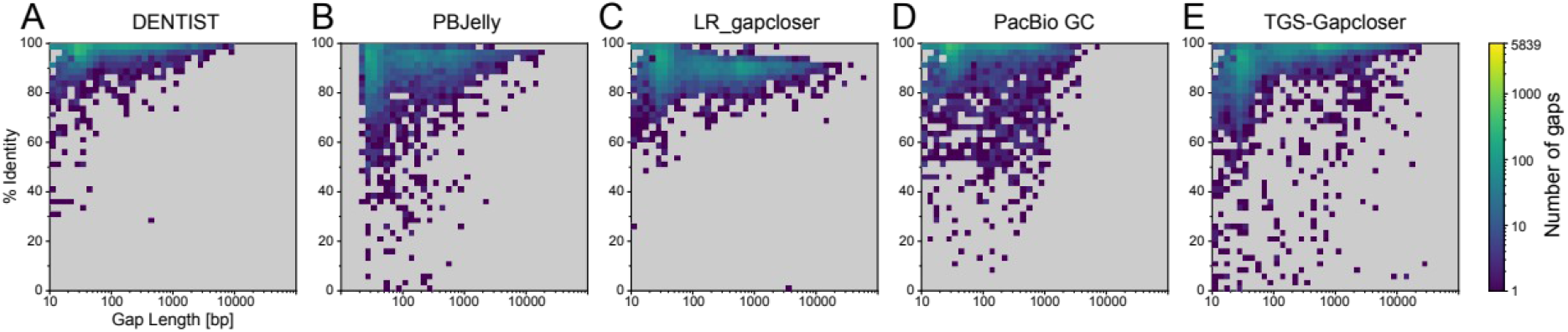
Comparison of base identity and gap size using the human assembly and real PacBio reads. (A) Gaps, which are closed at an accuracy of <90% by DENTIST, tend to be short. Many of these gaps contain simple repeats such as homopolymer runs, for which long reads have a higher error rate. In contrast, gaps closed at a lower accuracy by PBJelly (B), LR_gapcloser (C), PacBio GC (D) and TGS-Gapcloser (E) are more evenly distributed in size. For LR_gapcloser, there is a trend that the longer the gaps, the lower is the accuracy.

In terms of sensitivity, TGS-Gapcloser closed the most gaps for all assemblies except *Drosophila* where PacBio GC closed three additional gaps. TGS-Gapcloser achieved the highest contig NG50 increases for *Drosophila* and LR_gapcloser for the other three assemblies (Table 1, Figure 3D). DENTIST closed 74% (Human) to ∼95% (*Drosophila* and hummingbird) of the gaps, achieving the maximally possible contig NG50 for *Arabidopsis* and otherwise increasing NG50 7.5-fold (human) to 45.3-fold (hummingbird). Thus, consistent with the simulated read data, DENTIST has a reasonable sensitivity and the highest accuracy also when using real reads.

### Runtime and Memory Consumption

While a high accuracy and contiguity are certainly the most important objectives, speed and memory consumption of a gap closing method determine how large the required computational resources must be. We used the human dataset with real PacBio reads to compare the total CPU time and the maximum memory usage (measured as maximum resident set size across all jobs) of the gap closing tools. As shown in Table 2, TGS-Gapcloser is by far the fastest method (15.4 CPU hours), followed by DENTIST (255.5 CPU hours). TGS-Gapcloser required a slightly smaller amount of memory than DENTIST (24.6 Gb vs. 25.7 Gb). All other methods require significantly more memory and runtime. With this speed and memory consumption, DENTIST can finish a mammalian genome like human within a day on a 16-core workstation and within a few hours on a compute cluster, giving it also a good turn-around time for testing.

**Table 2:** Comparison of runtime and maximum memory consumption on the human assembly with real PacBio reads. For DENTIST, we did not count the runtime of local Snakemake rules, which consume very little resources. For PacBio GC, we excluded resources needed for the expensive pre- and post-processing of the assembly, but included the essential calls of *samtools faidx, pbindex, pbalign* and *variantCaller*. GiB = Gibibyte (1,073,741,824 = 2^30^ bytes).

| Tool | Total CPU Time [h] | Max. memory consumption [GiB] |
| --- | --- | --- |
| DENTIST | 255.5 | 25.7 |
| LR_gapcloser | 289.4 | 96.8 |
| PacBio GC | 5,904.2 | 40.6 |
| PBJelly | 2.285.0 | 53.9 |
| TGS-Gapcloser | **15.4** | **24.6** |

### Read Coverage Analysis

Another relevant consideration is how much coverage in long reads needs to be generated to close assembly gaps and thus improve an existing fragmented assembly. A higher read coverage likely allows one to close more gaps, but also increases the sequencing costs. To evaluate how DENTIST’s performance is affected by coverage, we ran our method with varying coverage of simulated PacBio reads on the *D*. *melanogaster* assembly test case. As shown in Figure 6, the number of closed gaps starts to plateau above a read coverage of about 15X. Also, above 15X the majority of gaps are closed with an accuracy ≥99%. Higher coverages increase the percent of gaps closed with ≥99% accuracy, since more reads facilitate the construction of a highly accurate consensus sequence. In summary, while assembly improvements are also possible with low coverages of 5X or 10X, a coverage between 15X and 20X appears to be a good tradeoff between sequencing cost and power to accurately close most assembly gaps. Importantly, this coverage is substantially less than the recommended coverage of ∼60X that is typically required for *de novo* assembly, providing a cost-effective alternative.

**Figure 6:**
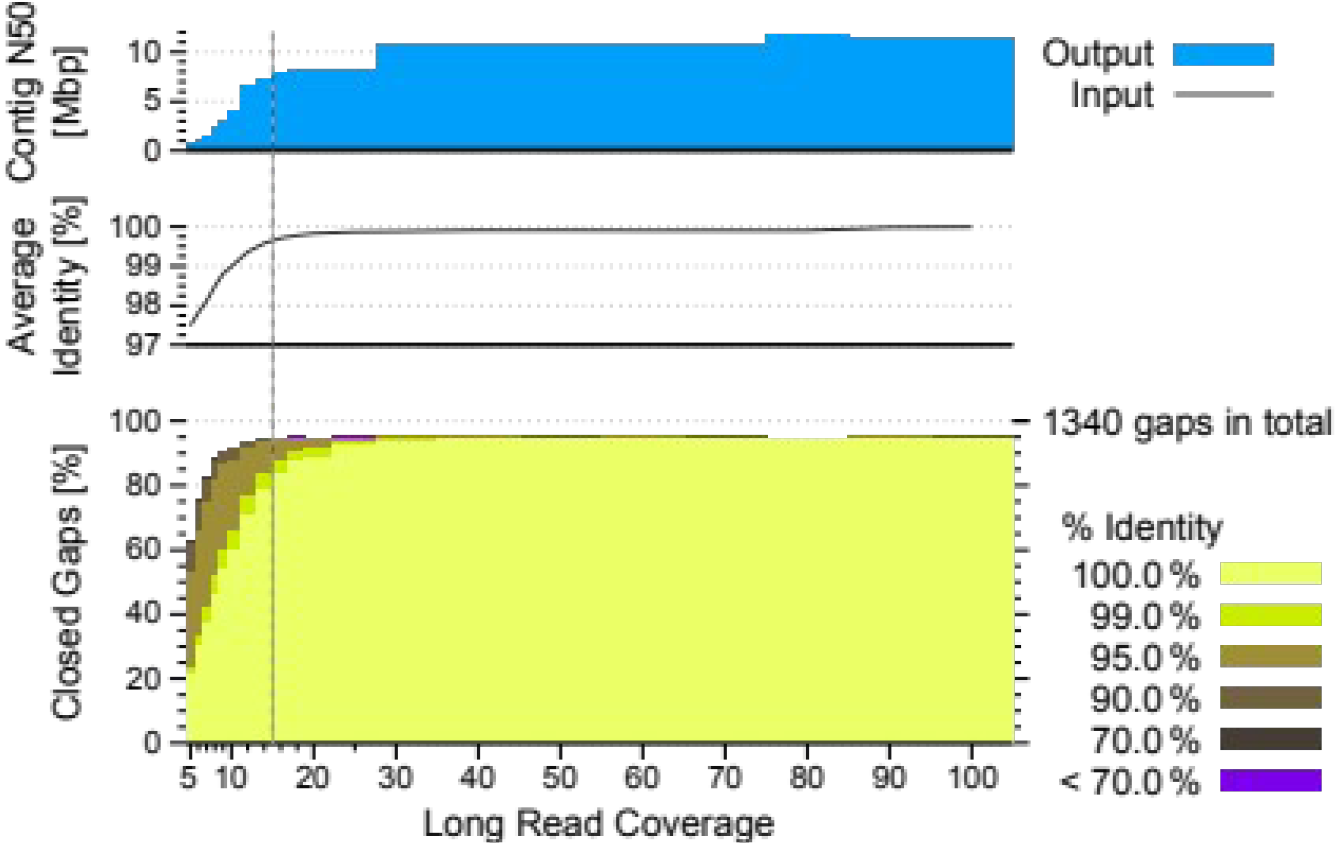
Coverage analysis by applying DENTIST to *Drosophila* with varying coverage of simulated PacBio CLR reads. The figure compares contig NG50 (top), average identity of the closed gaps (middle) and the of number of closed gaps together with a breakdown of their sequence identity (bottom). The number of closed gaps starts to plateau above a read coverage of about 15X (indicated by a vertical dashed line), which is significantly less than the recommended coverage of ∼60X required for *de novo* genome assembly.

## Discussion

We have presented a novel method, DENTIST, that uses uncorrected, long sequencing reads to close gaps in fragmented assemblies. DENTIST was developed with the main goal of closing assembly gaps at a very high accuracy, for which it implements a repeat-aware read alignment step to map reads to the correct assembly loci, a consensus sequence step to obtain an accurate sequence to fill a gap, and a final validation step. Our tests using simulated and real PacBio long-read data show that our method is substantially more accurate than existing tools, while achieving good sensitivity. Furthermore, DENTIST is sufficiently fast and memory-efficient to close gaps in a reasonable amount of time also in larger assemblies such as human. Together, these features make DENTIST appropriate for the task of improving the quality of hundreds of existing draft genomes with auxiliary long-read data.

The accuracy of a gap closing method is mainly influenced by the accuracy of the read mappings and by the ability to determine an accurate consensus from the error-prone long sequencing reads. The latter aspect is generally challenging and even with high read coverage it is difficult to reach a desired base accuracy of Q40 (99.99% base accuracy) [1]. Therefore, *de novo* genome assembly from long reads includes a final “polishing” step that maps shorter Illumina reads to the finalized assembly to correct remaining base errors [1, 2]. While DENTIST already achieves a high base accuracy of the sequence inserted into gaps, this accuracy is lower than Q40. Therefore, we recommend to polish the gap-closed assembly using Illumina reads after applying DENTIST. Notably, by achieving a base accuracy of 99% or higher, DENTIST facilitates the mapping of shorter Illumina reads, which would be more difficult with the lower accuracies produced by other methods.

To make it easy for users to run DENTIST, we used Snakemake [13] to automate the entire workflow (Figure 1). This pipeline is built in a modularized manner and is therefore customizable. Furthermore, to enable easy application on a compute cluster without the necessity of complicated software installation steps, we provide DENTIST and all required tools in a Docker container environment that can be easily used with Snakemake’s Singularity integration [14]. A conda package is also available. The full source code is available at https://github.com/a-ludi/dentist.

## Methods

### DENTIST parameters

The gap closing pipeline implemented in DENTIST requires a number of parameters that we empirically optimized with the goal of achieving a very high accuracy on the different test assemblies. All default parameter settings and how they can be adjusted by users via command line parameters are listed in the Supplementary Material (Listing 1, Note 1).

### Repeat masking and read mapping

Before aligning reads, DENTIST produces four types of repeat masks. First, it starts by masking low complexity regions in the given assembly using *DBdust* (https://dazzlerblog.wordpress.com/command-guides/dazz_db-command-guide/) in order to improve the sensitivity of the *daligner* alignment algorithm [15]. Second, tandem repeats are identified with *datander* and *TANmask* of the DAMASKER suite (https://dazzlerblog.wordpress.com/command-guides/damasker-commands/). Third, to identify other repetitive regions, DENTIST performs a self-alignment of the given assembly using *daligner* [15] and masks regions covered by at least four alignments to other genomic regions (adjustable via DENTIST’s parameter --max-coverage-self). Fourth, DENTIST creates another repeat annotation by analyzing the coverage of read alignments and marks assembly regions as repetitive that are covered by more reads than expected from the global read coverage (summed length of all long reads divided by genome size). To this end, DENTIST uses the first three repeat annotations as a soft mask and aligns all input long reads to the assembly using *damapper* (https://dazzlerblog.wordpress.com/2016/07/31/damapper-mapping-your-reads/), which outputs chains of local alignments arising from read artifacts such as poor quality regions or larger haplotype variations. All genomic regions covered by more than *C*_*max*_ alignments are considered repetitive. The threshold *C*_*max*_ is either given by the user via --max-coverage-reads or *C*_*max*_ is calculated from the global read coverage *C* (provided via --read-coverage) such that the probability of observing more than *C*_*max*_ alignments in a unique (non-repetitive) genomic region is very small (Supplementary Table 2). This probability is calculated under the assumption that the reads are sampled uniformly across the genome, implying a Poisson distribution of the number of sampled reads at any position in the genome (probability to observe *k* reads at any position is 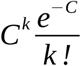). Genomic regions with a read coverage higher than *C*_*max*_ comprise the first part of the fourth (read alignment-based) repeat annotation. To further increase the sensitivity, DENTIST searches for smaller repeat-induced local alignments. To this end, we define an alignment as *proper* if there are at most 100 bp (adjustable via --proper-alignment-allowance) of unaligned sequence on either end of the read. All other alignments, where only a smaller substring of the read aligns, are called *improper*. Improper alignments are often indicative of repetitive regions. For example, an interspersed repeat inside a long read will result in improper alignments to other genomic loci, where a similar repeat copy exists. Therefore, DENTIST considers genomic regions where the number of improper read alignments is higher than a threshold to be repetitive. By default, this threshold equals half the global read coverage *C* (adjustable via --max-improper-coverage-reads). These genomic regions comprise the second part of the fourth repeat annotation.

These four repeat annotations are homogenized by transferring the annotated repeat regions to the reads using the read alignment and back again to the assembly. These homogenized, final repeat annotations are used in the next step of the pipeline.

### Alignment filtering

To extract candidate reads that could close assembly gaps from the entire set of read alignments, it is crucial to remove potentially unreliable and irrelevant alignments. A read alignment is categorized as reliable if (1) it is proper (defined above), (2) it is strongly anchored, i.e. at least 500 bp (adjustable via --min-anchor-length) of the aligned reference sequence are non-repetitive according to the homogenized repeat masks, and (3) after filtering improper alignments, every region of the aligned read must align at most to one assembly region. Finally, (4) alignments that are fully contained in a reference contig are removed because they are irrelevant for gap closing.

### Identifying closable gaps

DENTIST identifies closable gaps by creating a so-called *scaffold graph* that connects input contigs based on the filtered alignments of the long reads. For every contig in the assembly, the graph has four nodes v^i^_pre_, v^i^_begin_, v^i^_end_, and v^i^_post_representing virtual locations relative to the contig. As illustrated in Supplementary Figure 1, edges in this graph are interpreted as follows:

- {v^i^_begin_, v^i^_end_} represents contig i,
- {v^i^_end_, v^j^_begin_} for i≠j represent either reads that span contig *i* and *j* and thus also span the assembly gap between them, or intra-scaffold gaps (gaps between adjacent contigs) in the input assembly, and
- {v^i^_pre_, v^i^_begin_} and {v^i^_end_, v^i^_post_} represent reads extending the beginning or end of a contig, respectively.

Initially, the scaffold graph is populated with the contig and intra-scaffold gap edges. Then, for every read, edges of the types described above are inferred from its alignments. Note that for long reads that align to more than two consecutive contigs, we do not add the transitive edges (e.g. connecting contig i to i+2). A list of the reads and corresponding alignments is kept for every edge such that they can be retrieved for gap closing.

### Resolving scaffolding conflicts

The raw scaffold graph likely still contains some artifacts that need to be cleaned up. Scaffolding conflicts show up as begin nodes with more than one incoming edge or end nodes with more than one outgoing edge, meaning there is more than one way to connect the respective contig.

For assemblies with small contigs, a common conflict is small cycles resulting from reads that are long enough to completely cover one or more contigs and reach into both neighboring contigs. Depending on the local accuracy of the read, the repeat content of the intermediate contigs, and the minimum alignment length, local alignments to intermediate, small contigs may not be reliably detected, resulting in an edge that connects the neighboring contigs but skips the intermediate one(s). However, for other reads with higher accuracy, local alignments to these small intermediate contigs can be detected, resulting in a small cycle in the scaffolding graph. To avoid mistaking these cycles for scaffolding conflicts, DENTIST searches for small cycles (up to three intermediate contigs) in the scaffold graph, identifies the “skipping” edge and aligns the reads from that edge to the intermediate (skipped) contigs with increased alignment sensitivity. If this additional step detects an alignment between the skipping read and the intermediate contig(s), the new alignments will be used to correct the graph; otherwise the read will be discarded avoiding scaffolding conflicts. This procedure effectively resolves many branching points in the scaffold graph, while preserving valuable sequence information.

The remaining scaffolding conflicts are resolved by a crude yet effective heuristic: if there are two or more spanning edges at the begin or end node of a contig, then the spanning edge with the highest number of reads will be kept, if this edge is supported by at least three times (adjustable via --best-pile-up-margin) as many reads as for the other edges; otherwise all edges in question are discarded. Additionally, to use the given scaffolding information (contig order), edges that are supported by an intra-scaffold gap in the input assembly are given a higher weight by multiplying their read number with a bonus factor (default 6, adjustable via --existing-gap-bonus).

After removing scaffolding conflicts from the graph, DENTIST discards contig-spanning edges with fewer than three reads, as no accurate consensus sequence for the gap can be determined from one or two reads. Also, more spanning reads give higher confidence about the correctness of the join. The minimum number of spanning reads can be increased via the parameter --min-spanning-reads to achieve a higher confidence. DENTIST also provides an “expert option” to allow gap-closing with solitary reads in which case the raw read sequence would be inserted. For valid contig-spanning edges, DENTIST puts the reads representing the spanning and extending edges into a single pile up to gather as much sequence information as possible for the consensus procedure.

### Closing the gaps

To compute a consensus sequence for a gap, reads assigned to spanning or extending edges are cropped such that all read alignments begin and end at the same position in the reference assembly. The cropping position is chosen such that all reads still overlap the flanking contigs as much as possible to allow validation of the consensus. Cropping reduces artifacts in the subsequent pairwise alignment of the involved reads and allows easy identification of false alignments.

The cropped reads are pairwise aligned to each other using *daligner*. These read-to-read alignments often contain several local alignments, because long reads can contain regions of poor quality or larger indels and because complex gaps can result in local repeat-induced alignments. Before computing a consensus sequence, these local alignments must be filtered and “chained”. To this end, we implemented a chaining algorithm that works directly with the alignment output produced by *daligner*. This algorithm is applied to every read-to-read alignment and reduces the alignment chaining problem to the shortest paths problem on a directed acyclic graph with node and edge weights. Every local alignment is represented by a node with a negative (beneficial) weight corresponding to the average number of base pairs covered by the local alignment of the involved reads. To determine edge weights, we define *gap*_*A*_*(x,y)andgap*_*B*_*(x,y)*as the distance between two ordered local alignments *x*and *y*on long reads *A* and *B*, respectively, and *gapSizeDiff(x,y)*as the absolute difference between *gap*_*A*_*(x,y)*and *gap*_*B*_*(x,y)*. Thus, *gap*_*A*_*(x,y)*represents the number of unaligning bases in read A between *x* and *y*, which is 0 if both local alignments are adjacent, and *gapSizeDiff(x,y)*represents the difference in the number of unaligned bases in both reads. Two nodes in the graph are connected by an edge if the respective local alignments *x*and *y*are *chainable*, i.e. (1) the first alignment begins strictly before the second alignment and both occur in the same orientation, (2) *gapSizeDiff(x,y)*< 1000 bp (adjustable via parameter --max-indel), (3) *gap*_*A*_*(x,y)*and *gap*_*B*_*(x,y)*are smaller than 10,000 bp (adjustable via --max-chain-gap), and (4) the relative overlap between the alignments, determined as the length of the overlapping region divided by the length of the smaller alignment, is ≤0.3 (adjustable via --max-relative-overlap). The edge then gets a positive weight, penalizing the difference between the number of unaligning bases and to smaller extent the length of the unaligning region between both local alignments as *w*_*e*_*(x,y)*= *gapSizeDiff(x,y)*+ 0.1max·{| *gap*_*A*_*(x,y)*|, |*gap*_*B*_*(x,y)*|}. Maximal shortest paths in this graph constitute candidates for alignment chains. From all candidate chains, the best scoring chain(s) are selected.

After chaining, intrinsic quality values (QVs) are derived from the alignment chains using *DAScover*and *DASqv*(https://dazzlerblog.wordpress.com/command-guides/dascrubber-command-guide/). The read with the lowest number of bad QVs is chosen as the reference read, where a QV is defined as bad if it belongs to the 8% worst QVs in the pileup (adjustable via --bad-fraction). The cleaned up alignment is used as input for *daccord* [16], which uses a local de Bruijn graph-based approach to compute an accurate consensus based on the selected reference read.

Subsequently, the consensus sequence will be aligned to the flanking contigs to validate its correctness and find the exact insert sites such that the contigs are not modified when closing the gap with the consensus sequence. DENTIST allows closing gaps in three different modes. In the default mode, DENTIST closes only intra-scaffold gaps that are provided in the input assembly. In the second mode, DENTIST additionally closes gaps between different scaffolds if they are spanned by sufficient long read evidence (minimum of three spanning reads by default). In this mode, both intra- and inter-scaffold gaps are closed. In the third mode, DENTIST uses the existing scaffolding information (if available) only for conflict resolution and freely scaffolds the given contigs using the long reads. The second and third mode enables DENTIST to also improve contig-only assemblies. Note that the selection of candidates for gap closing is only based on the evidence derived from the input read data and does not depend on the chosen mode.

### Validation of closed gaps

Aiming at a high accuracy, DENTIST performs a final validation step by mapping the input reads to the gap-closed assembly. For each closed gap, DENTIST analyzes the genomic region 1000 bp (adjustable via --region-context) up- and downstream of the former gap. A closed gap is validated if there are at least three (adjustable via --min-spanning-reads) unchained read alignments spanning this region and the minimum *continuous alignment coverage* exceeds a user-given threshold, definable via the mandatory parameter --min-coverage-reads (alternatively, if the user provides the global long-read coverage via --read- coverage and the ploidy via --ploidy, DENTIST will set the threshold to 50% of the long-read coverage expected to be sequenced from a haploid locus). The *continuous alignment coverage* with window size *w* (default 500 bp; adjustable via --weak-coverage-window) at position *x* is defined as the number of local alignments (unchained and potentially improper) that completely cover the window [*x, x*+ *w*). The minimum *continuous alignment coverage* is then obtained by sliding the window across all positions in the genomic region defined above. For closed gaps that are not validated, DENTIST outputs the original gap (NNN…) sequence.

### Evaluating the gap closing accuracy with realistic assembly gap sizes and loci

To assess the accuracy of DENTIST and compare it to other methods, we devised a realistic ground truth scenario, where the true sequence of assembly gaps is considered to be known and where assembly gaps occur at realistic genomic loci and at realistic sizes. This is important as repetitive genomic regions are the main reason for assembly gaps, thus random assembly gap placement or replacing repeats with gaps will not create a realistic setting. Each test scenario comprises (1) a high-quality reference assembly, which we consider as the ground truth, (2) a test assembly that contains assembly gaps and is input to the gap closing method, and (3) a set of long reads that were either simulated using the reference assembly or are real PacBio reads (Supplementary Table 1). Subsequently, we derived performance statistics from the output assembly of a gap closer using the same evaluation strategy.

To generate a test assembly with realistic gap sizes and loci, we aligned the reference assembly to a more fragmented, short-read assembly of the same species and copied gaps from short-read assembly into the respective position of the reference assembly, as depicted in Figure 2. Briefly, we aligned both assemblies using *lastz* [17] and constructed *liftOver* chains [18]. Then, we obtained the 500 bp upstream and downstream flanks of each gap in the short-read assembly and used *liftOver* [19] (parameter -minMatch=0.8) to map these flanks to the reference assembly. Before introducing assembly gaps into the reference, we disassembled the reference assembly into its contigs to avoid having gaps without a known ground truth sequence. For a pair of flanks that belong to the same gap and were mapped adjacently to a reference contig, we replaced the real sequence (ground truth) between both flanks with N’s with *dentist build-partial-assembly*, if the gap size is at least 10 bp. We required that the reference assembly itself (which is a real assembly with gaps on its own) does not already have a gap within 3 kb of an introduced gap. It should be noted that the introduced gap size in our tests is identical to the size of the ground truth gap sequence. This gives gap closers that take gap size into account an advantage (e.g. PBJelly, LR_gapcloser and PacBio GC) but not DENTIST, which does not use the estimated size of gaps as in reality they are often imprecise or even have a pseudo-size not related to the real gap size at all.

Since the true sequence in gaps is known, it is possible to automatically assess the performance of the gap closing tools. To this end, we first identify the original contigs of the test assembly in a gap-closed assembly by searching for exact and unique matches. Duplicated contigs that already have more than one exact match within the test assembly are excluded. We noticed that some gap closing tools do not only fill in gaps but also modify the input contigs. We handled this unexpected behavior in two ways. First, before searching for exact matches, we remove a given number of base pairs from the flanks of the contig of the test assembly, thus allowing the gap closer to modify contig flanks. This number of base pairs in controlled by --crop-ambiguous (default 100bp) when searching for duplicate contigs and by --crop-alignment when searching for contigs in the gap-closed assembly. Second, after identifying exact matches, we optionally conduct a second search for the remaining contigs, allowing for up to 1.5% mismatches but requiring that a contig fully and uniquely aligns to some part of the result assembly. Consequently, original contigs that are substantially modified by a gap closing method may not be detected, which highlights an unwanted behavior of the respective method.

After determining the locations of the test contigs, each gap is evaluated according to the following rules. A gap is considered *closed* if the locations of both contigs surrounding the gap in the test assembly are mapped to a single contig in the gap-closed assembly. In this case, the sequence identity between the known gap sequence and the inserted gap sequence is calculated. A gap is considered *unclosed* if the locations of both contigs surrounding the gap in the test assembly are known and the contigs are mapped to different but adjacent contigs in the gap-closed assembly, i.e. a gap remains. A gap is considered *unknown* if none of the flanking contigs are found or at least one flanking contigs cannot be located uniquely. These gaps are excluded from further analysis, as we cannot accurately determine the inserted gap sequence. If neither of the above cases is true, the gap is said to be *broken*. This is the case if both flanking contigs could be located but are not adjacent anymore, i.e. the contigs are misassembled. This procedure allows to obtain an accurate assessment of the performance of each gap closing tool in an automated fashion.

Long reads used in our tests were either real PacBio reads (Supplementary Table 1) or reads that were sampled (simulated) from the reference assembly adding a typical PacBio base and indel error profile using “*simulator-m25000-s125000-e.13-rSEED”*(https://dazzlerblog.wordpress.com/command-guides/dazz_db-command-guide/). These parameters provide a length and error rate distribution that match the distributions of current CLR reads. The seed used for simulating reads is listed in Supplementary Table 1. For real PacBio reads, we provided all subreads from all wells to the gap closing tools.

### Running gap closing methods

All tools were called with the default or recommended parameters, except for parameters that do not influence the output such as the number of parallel threads. Supplementary Table 3 lists the exact parameters used for each tool. For *finisherSC.py*, we used parameters “-- fast True --large True” to make the *Drosophila* application feasible in terms of runtime and memory requirements. For PacBio GC, we ran *variantCaller* with parameter “-- algorithm=best”, which automatically selects Arrow or Quiver as the most appropriate method. To prevent PacBio GC from modifying bases in the contigs outside of gaps, which hampers the exact identification of the inserted sequences, we restricted its application to the exact gap locations using the “--referenceWindowsFile” parameter. Furthermore, we distributed the computation on our compute cluster by (1) computing the read alignments for the whole assembly, (2) splitting the assembly into blocks of ∼200 Mb and dividing the read alignments accordingly, (3) applying PacBio GC to each block separately, and (4) merging the unmodified contigs with the processed gaps into the output assembly. This complete workflow can be found at https://bds.mpi-cbg.de/hillerlab/DENTIST/source/arrow/Snakefile.

The results were evaluated using “*dentist check-results* ” with parameter “--crop-ambiguous=300” to ignore 300 bp from the contig flanks when searching for duplicate contigs. To evaluate PBJelly, parameter “--crop-alignment=100” was used in addition to ignore 100 bp from the contig flanks in the contig identification step. To evaluate LR_gapcloser, which can modify not only the original contig flanks, we additionally specified parameters “--crop-alignment=300” and “--recover-imperfect-contigs” to enable contig identification with up to 1.5% mismatches. The increase in contiguity of the gap-closed assembly was measured by the percentage of closed gaps and the increase in contig NG50. The correctness of closed gap sequences was measured as the sequence identity between the (known) ground truth and the inserted sequence for each gap. These values are presented in three forms: as a distribution binned in six intervals: [0, 0.7), [0.7, 0.9), [0.9, 0.95), [0.95, 0.99), [0.99, 1.0), and {1.0}, as the arithmetic mean over all the sequence identities, and as the weighted arithmetic mean using the true gap sizes as weights.

For tests with simulated reads, we also used “*dentist find-closable-gaps”* to determine which gaps are *closable*, as the true origin of each read is known. We defined a gap as *closable* if and only if at least three reads span the gap and 500 bp on either side.

## Supporting information

Supplementary Figures & Remarks

## Data availability

All data, including the reference and test assemblies with introduced gaps and their true sequence as valuable data for future method comparisons, is available at https://bds.mpi-cbg.de/hillerlab/DENTIST/.

## Availability of supporting source code and requirements

Project name: DENTIST

Project home page: https://github.com/a-ludi/dentist

Operating systems: Linux

Programming language: D

Other requirements: Snakemake 5.32.1 or higher

License: MIT

RRID: SCR_021856

biotools: dentist

## Author contributions

AL implemented the software and analyzed the data. MP tested the software. MH and GM conceived and supervised this study. MH and AL wrote the manuscript. All authors read and approved the final manuscript.

## Acknowledgment

We thank the Computer Service Facilities of the MPI-CBG for their support.

