## Supplementary Figures & Remarks for "DENTIST – using long reads for closing assembly gaps at high accuracy"

Supplementary Information contains Supplementary Figure 1, Listing 1 and Note 1. Supplementary Tables 1-3 are provided in a separate Excel file.

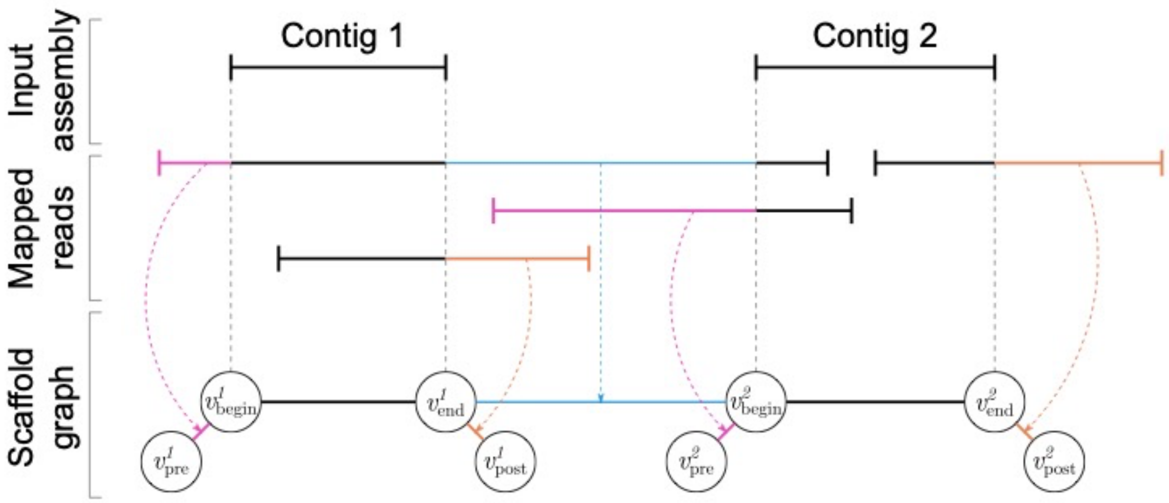


**Supplementary Figure 1:** Visualization of a scaffold graph for two exemplary contigs.

For each contig, four nodes $v_{pre}^i$, $v_{begin}^i$, $v_{end}^i$ and $v_{post}^i$ exist. Edges between nodes represent the contig itself (black), parts of a reads that span and thus connect contigs (blue) or parts of reads that extend a contig (purple and orange).

### Listing 1: List of command line options for DENTIST

The following list contains all command line options of DENTIST. Default values are given in parentheses after the option name and the associated commands are stated in parentheses after the colon.

**Example:**

```
--fasta-line-width, -w <ulong>(50): (output)
                              └──┘   └──────┘
                       default value    command
```

**List of Options:**

- --agp <string>: (output)
  write AGP v2.1 file that describes the output assembly
- --allow-single-reads: (process-pile-ups)
  allow using single reads instead of consensus sequence for gap closing
- --auxiliary-threads, --aux-threads, -A num-threads(floor(totalCpus / <threads>): (collect-pile-ups, process-pile-ups)
  use <num-threads> threads for auxiliary tools like daligner, damapper and daccord)
- --bad-fraction <frac>(0.8): (process-pile-ups)
  Intrinsic QVs are categorized as “bad” if they are greater or equal to the best QV of the worst <frac> trace point intervals.
- --batch, -b <idx-spec>[,<idx-spec>...]: (process-pile-ups)
  process only a subset of the pile ups. <pile-up-ids> is a comma-separated list of <idx-spec>. Each <id-specifications> is either a single integer <idx> or a range <from>..<to>. <idx>, <from> and <to> are zero-based indices into the pile up DB. The range is right-open, i.e. index <to> is excluded. <to> may be a dollar-sign ($) to indicate the end of the pile up DB.
- --bed <string>(standard input): (bed2mask)
  input BED file; fields must be TAB-delimited
- --best-pile-up-margin <double>(3.0): (collect-pile-ups)
  given a set of conflicting gap closing candidates, if the largest has <double> times more reads than the second largest it is considered unique. If a candidates would close gap in the reference assembly marked by ns the number reads is multiplied by –existing-gap-bonus.
- --cache-contig-alignments <string>( ): (validate-config)
  if given the contig location will be cached as JSON faking the effect of the same option in check-results. NOTE: the result has to amended manually to be fully valid.
- --closed-gaps-bed <string>: (output)
  write BED file with coordinates of closed gaps
- --config <config-json>: (all except validate-config)
  provide configuration values in a JSON file. See README.md for usage and examples.

- --daccord <daccord-option>[,<daccord-option>...]: (process-pile-ups)
  Provide additional options to daccord
- --daligner-consensus <daligner-option>[,<daligner-option>...]: (process-pile-ups)
  Provide additional options to daligner
- --daligner-reads-vs-reads <daligner-option>[,<daligner-option>...]: (process-pile-ups)
  Provide additional options to daligner
- --daligner-self <daligner-option>...: (generate-dazzler-options, process-pile-ups)
  Provide additional options to daligner
- --damapper-ref-vs-reads <damapper-option>...: (generate-dazzler-options, collect-pile-ups)
  Provide additional options to damapper
- --data-comments: (bed2mask)
  parse BED comments (column 4) as generated by output. This will cause a crash if formatting errors are encountered.
- --datander-ref <datander-option>[,<datander-option>...]: (generate-dazzler-options, process-pile-ups)
  Provide additional options to datander
- --debug-pile-ups <db-stem>: (collect-pile-ups)
  write pile ups of intermediate steps to <db-stem>.<state>.db
- --debug-repeat-masks: (mask-repetitive-regions)
  (only for reads-mask) write mask components into additional masks <repeat-mask>-<component-type>
- --dust-reads <dust-option>[,<dust-option>...]: (process-pile-ups)
  Provide additional options to dust
- --dust-ref <dust-option>[,<dust-option>...]: (generate-dazzler-options)
  Provide additional options to dust
- --existing-gap-bonus <double>(6.0): (collect-pile-ups)
  if a candidate would close an existing gap its size is multiplied by <double> before conflict resolution (see –best-pile-up-margin).
- --fasta-line-width, -w <ulong>(50): (output)
  line width for output FASTA
- --help, -h: (all)
  Prints this help.
- --join-policy <JoinPolicy>(scaffoldGaps): (output)
  allow only joins (gap filling) in the given mode: scaffoldGaps (only join gaps inside of scaffolds – marked by ns in FASTA), scaffolds (join gaps inside of scaffolds and try to join scaffolds), contigs (break input into contigs and re-scaffold everything; maintains scaffold gaps where new scaffolds are consistent)

- `--json, -j`: (`show-mask`, `show-pile-ups`, `show-insertions`, `translate-coords`)
  if given write the information in JSON format
- `--keep-temp, -k`: (`collect-pile-ups`, `process-pile-ups`)
  keep the temporary files; outputs the exact location
- `--mask, -m <name>[,<name>...]`: (`propagate-mask`, `collect-pile-ups`, `process-pile-ups`)
  Dazzler masks for repetitive regions (at least one required; generate with `mask-repetitive-regions` command)
- `--max-chain-gap <bps>(10000)`: (`chain-local-alignments`, `process-pile-ups`)
  two local alignments may only be chained if at most <bps> of sequence in the A-read and B-read are unaligned.
- `--max-coverage-reads <uint>`: (`mask-repetitive-regions`)
  this is used to derive a repeat mask from the ref vs. reads alignment; if the alignment coverage is larger than <uint> it will be considered repetitive; a default value is derived from –read-coverage; both options are mutually exclusive
- `--max-coverage-self <uint>(4)`: (`mask-repetitive-regions`)
  this is used to derive a repeat mask from the self-alignment; if the alignment coverage larger than <uint> it will be considered repetitive
- `--max-improper-coverage-reads <uint>`: (`mask-repetitive-regions`)
  this is used to derive a repeat mask from the ref vs. reads alignment; if the coverage of improper alignments is larger than <uint> it will be considered repetitive; a default value is derived from –read-coverage; both options are mutually exclusive
- `--max-indel <bps>(1000)`: (`chain-local-alignments`, `process-pile-ups`)
  two local alignments may only be chained if the resulting insertion or deletion is at most <bps>
- `--max-insertion-error <double>(0.10)`: (`output`)
  insertion and existing contigs must match with less error than <double>
- `--max-relative-overlap <fraction>(0.30)`: (`chain-local-alignments`, `process-pile-ups`)
  two local alignments may only be chained if the overlap between them is at most <fraction> times the size of the shorter local alignment. This must hold for the reference and query.
- `--min-anchor-length <uint>(500)`: (`generate-dazzler-options`, `collect-pile-ups`, `process-pile-ups`)
  alignment need to have at least this length of unique anchoring sequence
- `--min-coverage-reads <num>`: (`validate-regions`)
  validly closed gaps must have a continuous coverage of at least <num> properly aligned reads; see –weak-coverage-mask for more details
- `--min-extension-length <ulong>(100)`: (`output`)
  extensions must have at least <ulong> bps of consensus to be inserted
- `--min-gap-size <uint>(0)`: (`filter-mask`)
  minimum size for gaps between mask intervals

- `--min-interval-size <uint>(0)`: (`filter-mask`)
  minimum size for mask intervals
- `--min-reads-per-pile-up <ulong>(3)`: (`process-pile-ups`)
  pile ups must have at least <ulong> reads to be processed
- `--min-relative-score <fraction>(1.0)`: (`chain-local-alignments`, `process-pile-ups`)
  output chains with a score of at least <fraction> of the best chains score. A value of 1.0 means that only chains with the best chains score will be accepted; a value of 0.0 means that all chains will be accepted
- `--min-score <int>(trace point spacing of alignment)`: (`chain-local-alignments`, `process-pile-ups`)
  output chains with a score of at least <int>
- `--min-spanning-reads, -s <ulong>(3)`: (`collect-pile-ups`, `validate-regions`)
  require at least <ulong> spanning reads to close a gap
- `--no-highlight-insertions, -H`: (`output`)
  turn off highlighting (upper case) of inserted sequences in the FASTA output
- `--no-merge-extension`: (`collect-pile-ups`)
  Do not merge extension reads into spanning pile ups.
- `--only <OnlyFlag>(spanning)`: (`process-pile-ups`, `output`)
  only process/output insertions of the given type. Note, extending insertions are experimental and may produce invalid results.
- `--ploidy, -N <uint>`: (`validate-regions`)
  this is used to derive a lower bound for the read coverage
- `--progress`: (`chain-local-alignments`)
  Print regular status reports on the progress.
- `--progress-every <msecs>(500)`: (`chain-local-alignments`)
  Print status reports every <msecs>.
- `--progress-format <format>(human)`: (`chain-local-alignments`)
  Use <format> for status report lines where <format> is either `human` or `json`. The former prints a status line that updates regularly while the latter prints a full JSON record per line with every update
- `--proper-alignment-allowance num(trace point spacing of alignment)`: (`mask-repetitive-regions`, `collect-pile-ups`, `process-pile-ups`, `validate-regions`)
  An alignment is called proper if it is end-to-end with at most <num> bp allowance.
- `--quiet, -q`: (all)
  reduce output as much as possible reporting only fatal errors. If given this option overrides –verbose.
- `--read-coverage, -C <double>`: (`mask-repetitive-regions`, `validate-regions`)
  this is used to provide good default values for –max-coverage-reads or –min-coverage-reads; both options are mutually exclusive
- `--region-context <bps>(1000)`: (`validate-regions`)
  consider <bps> base pairs of context for each region to detect splicing errors

- `--report-all`: (`validate-regions`)
  report all validation results instead of only failed gaps
- `--revert <option>[,<option>...]`: (all)
  revert named option to default value. This is useful to revert specific options of a config file.
- `--scaffolding <insertions-db>`: (`output`)
  write the assembly scaffold to <insertions-db>; use `show-insertions` to inspect the result
- `--skip-gaps <gap-spec>[,<gap-spec>...]`: (`output`)
  Do not close the specified gaps. Each <gap-spec> is a pair of contig IDs <contigA>-<contigA> meaning that the specified contigs should not be closed. They will still be joined by a preexisting gap.
- `--skip-gaps-file <file>`: (`output`)
  Same as –skip-gaps but <file> contains one <gap-spec> per line. If both options are given the union of all <gap-spec>s will be used. Empty lines and lines starting with `#` will be ignored.
- `--threads, -T <uint>(number of cores)`: (`collect-pile-ups`, `process-pile-ups`, `validate-regions`)
  use <uint> threads
- `--tmpdir, -P <string>`: (`collect-pile-ups`, `process-pile-ups`)
  use <string> as a working directory
- `--usage`: (all)
  Print a short command summary.
- `--verbose, -v`: (all)
  increase output to help identify problems; use up to three times. Warning: performance may be drastically reduced if using three times.
- `--weak-coverage-mask <mask>`: (`validate-regions`)
  write a Dazzler mask <mask> of weakly covered regions, i.e. sliding windows of –weak-coverage-window base pairs are spanned by less than –min-coverage-reads local alignments
- `--weak-coverage-window <bps>(500)`: (`validate-regions`)
  consider sliding window of <bps> base pairs to identify weak coverage

### Remark 1: Guide on configuration of DENTIST

DENTIST comprises a complex pipeline of with many options for tweaking. This section points out some important parameters and their effect on the result or performance.

The default parameters are rather **conservative**, i.e. they focus on correctness of the result while not sacrificing too much sensitivity.

We also provide a **greedy** sample configuration (`snakemake/dentist.greedy.json`) which focuses on sensitivity but may introduce more errors. **Warning:** *Use with care! Always validate the closed gaps (e.g. manual inspection).*

In any case, the workflow creates an intermediate assembly `workdir/{output_assembly}-preliminary.fasta` that contains all closed gaps, i.e. before validation. It is accompanied by an AGP and BED file. You may inspect these file for maximum sensitivity.

#### How to Choose DENTIST Parameters

While listing 1 is a good reference, it does not provide an overview of the important parameters. Therefore, we provide this shorter list of important and influential parameters. Please also consider adjusting the performance parameter in the workflow configuration (`snakemake/snakemake.yml`).

- `--dust-{reads,ref}`, `--daligner-{consensus,reads-vs-reads,self}`, `--damapper-ref-vs-reads`, `--datander-ref`, `--daccord`: These options allow passing parameters to the respective tools. They may have dramatic influence on the result. The default settings work well for PacBio CLR reads and should also work well with raw Nanopore data.

  In-depth discussion of each tool goes beyond the scope of this document, please refer to the respective documentations (DBdust, daligner, damapper, datander, daccord).

- `--max-insertion-error`: Strong influence on quality and sensitivity. Lower values lead to lower sensitivity but higher quality. The maximum recommended value is `0.05`.

- `--min-anchor-length`: Higher values results in higher accuracy but lower sensitivity. Especially, large gaps cannot be closed if the value is too high. Usually the value should be at least `500` and up to `10_000`.

- `--min-reads-per-pile-up`: Choosing higher values for the minimum number of reads drastically reduces sensitivity but has little effect on the quality. Small values may be chosen to get the maximum sensitivity in *de novo* assemblies. Make sure to throughly validate the results though.

- `--min-spanning-reads`: Higher values give more confidence on the correctness of closed gaps but reduce sensitivity. The value must be well below the expected coverage.

- `--allow-single-reads`: May be used under careful consideration in combination with `--min-spanning-reads=1`. This is intended for one of the following scenarios:

  1. DENTIST is meant to close as many gaps as possible in a *de novo* assembly. Then the closed gaps must be validated by other means afterwards.
  2. DENTIST is used not with real reads but with an independent assembly.

- `--existing-gap-bonus`: If DENTIST finds evidence to join two contigs that are already consecutive in the input assembly (i.e. joined by `N`s) then it will preferred over conflicting joins (if present) with this bonus. The default value is rather conservative, i.e. the preferred join almost always wins over other joins in case of a conflict.

- `--join-policy`: Choose according to your needs:
    1. `scaffoldGaps`: Closes only gaps that are marked by `N`s in the assembly. This is the default mode of operation. Use this if you do not want to alter the scaffolding of the assembly. See also `--existing-gap-bonus`.
    2. `scaffolds`: Allows whole scaffolds to be joined in addition to the effects of `scaffoldGaps`. Use this if you have (many) scaffolds that are not yet full chromosome-scale.
    3. `contigs`: Allows contigs to be rearranged freely. This is especially useful in *de novo* assemblies **before** applying any other scaffolding methods as it increases the contiguity thus increasing the chance that large-scale scaffolding (e.g. Bionano or Hi-C) finds proper joins.
- `--min-coverage-reads`, `--min-spanning-reads`, `--region-context`: DENTIST validates closed gaps by mapping the reads to the gap-closed assembly. It requires for each gap and the base pairs down- and upstream (`--region-context`) are (1) covered by at least `--min-coverage-reads` reads at every position and (2) are spanned by at least `--min-spanning-reads` reads. Thus, increasing any of these numbers makes the *valid* gaps more robust but may reduce their number.

#### Choosing the Read Type

In the examples PacBio long reads are assumed but DENTIST can be run using any kind of long reads. Currently, this is either PacBio or Oxford Nanopore reads. For using none-PacBio reads, the `reads_type` in `snakemake.yml` must be set to anything other than `PACBIO_SMRT`. The recommendation is to use `OXFORD_NANOPORE` for Oxford Nanopore. These names are borrowed from the NCBI. Further details on the rationale can found in this issue.

#### Cluster/Cloud Execution

Cluster job schedulers can become unresponsive or even crash if too many jobs with short running time are submitted to the cluster. It is therefore advisable to adjust the workflow accordingly. We tried to provide a default configuration that works in most cases as is but the application scenarios can be very diverse and manual adjustments may become necessary. Here is a small guide which config parameters influence the number of jobs and how much resources they consume.

- `max_threads`: Sets the maximum number of threads/cores a single job may use. A single-threaded job will always allocate a single core but thread-parallel steps, e.g. the sequence alignments, will use up to `max_threads` if snakemake has been provided enough cores via `--cores`.
- `-s<block_size:uint>`: The assembly and reads FAST/A files are converted into Dazzler DBs. These DBs store the sequence in a 2-bit encoding and have additional features like tracks (similar to BED files). Also they are split into blocks of `<block_size>`Mb. Alignments are calculated on the basis of these blocks which enables easy distribution onto the cluster. The larger the block size the longer are the alignment jobs and the more memory they require but also the number of jobs is reduced. Experience shows that the block size should be between 200Mb and 500Mb.
- `propagate_batch_size`: The repeat masks are homogenized by propagating them from the assembly to the reads and back again. Usually these jobs are very short because the propagation is parallelized over the blocks of the reads DB. To reduce the number of jobs both propagation directions are grouped together and submitted in

batches of propagate_batch_size read blocks. Increasing propagate_batch_size reduces the number of submitted jobs and increases the run time per job. It has no effect on the memory requirements.

- batch_size: In the collect step DENTIST identifies candidates for gap closing each consisting of a pile up of reads. From these pile ups consensus sequences are computed and validated in the process step. Each job process batch_size pile ups. Increasing batch_size reduces the number of submitted jobs and increases the run time per job. It has no effect on the memory requirements.
- validation_blocks: The preliminarily closed gaps are validated by analyzing how the reads align to each closed gap. The validation is conducted in independent jobs for validation_blocks many blocks of the gap-closed assembly. Decreasing validation_blocks reduces the number of submitted jobs and increases the run time and memory requirements per job. The memory requirement is proportional to the size of the read alignment blocks.
